# Larger GPU-accelerated brain simulations with procedural connectivity

**DOI:** 10.1101/2020.04.27.063693

**Authors:** James C Knight, Thomas Nowotny

## Abstract

Large-scale simulations of spiking neural network models are an important tool for improving our understanding of the dynamics and ultimately the function of brains. However, even small mammals such as mice have on the order of 1 × 10^12^ synaptic connections which, in simulations, are each typically charaterized by at least one floating-point value. This amounts to several terabytes of data – an unrealistic memory requirement for a single desktop machine. Large models are therefore typically simulated on distributed supercomputers which is costly and limits large-scale modelling to a few privileged research groups. In this work, we describe extensions to GeNN – our Graphical Processing Unit (GPU) accelerated spiking neural network simulator – that enable it to ‘procedurally’ generate connectivity and synaptic weights ‘on the go’ as spikes are triggered, instead of storing and retrieving them from memory. We find that GPUs are well-suited to this approach because of their raw computational power which, due to memory bandwidth limitations, is often under-utilised when simulating spiking neural networks. We demonstrate the value of our approach with a recent model of the Macaque visual cortex consisting of 4.13 × 10^6^ neurons and 24.2 × 10^9^ synapses. Using our new method, it can be simulated on a single GPU – a significant step forward in making large-scale brain modelling accessible to many more researchers. Our results match those obtained on a supercomputer and the simulation runs up to 35 % faster on a single high-end GPU than previously on over 1000 supercomputer nodes.

## 1. Introduction

The brain of a mouse has around 70 × 10^6^ neurons, but this number is dwarfed by the 1 × 10^12^ synapses which connect them [Herculano-Houzel et al., 2006]. In computer simulations of spiking neural networks, propagating spikes involves adding the synaptic input from each spiking presynaptic neuron to the postsynaptic neurons. The information describing which neurons are synaptically connected and with what weight is typically generated before a simulation is run and stored in large arrays. For large-scale brain models this creates high memory requirements, so that they can typically only be simulated on large distributed computer systems using software such as NEST [Gewaltig and Diesmann, 2007] or NEURON [Carnevale and Hines, 2006]. By careful design, these simulators can keep the memory requirements for each node constant, even when a simulation is distributed across thousands of nodes [Jordan et al., 2018]. However, high performance computer (HPC) systems are bulky, expensive and consume a lot of power and are hence typically shared resources, only accessible to a limited number of researchers and for time-limited investigations.

Neuromorphic systems [Frenkel et al., 2018, Furber et al., 2014, Merolla et al., 2014, Qiao et al., 2015, Schemmel et al., 2017] take inspiration from the brain and have been developed specifically for simulating large spiking neural networks more efficiently. One particular relevant feature of the brain is that its memory elements – the synapses – are co-located with the computing elements – the neurons. In neuromorphic systems, this often translates to dedicating a large proportion of each chip to memory. However, while such on-chip memory is fast, it can only be fabricated at relatively low density so that many of these systems economize – either by reducing the maximum number of synapses per neuron to as few as 256 or by reducing the precision of the synaptic weights to 6 [Schemmel et al., 2017], 4 [Frenkel et al., 2018] or even 1 bit [Merolla et al., 2014]. This allows some classes of spiking neural networks to be simulated very efficiently, but reducing the degree of connectivity to fit within the constraints of current neuromorphic systems inevitably changes the dynamics of brain simulations [van Albada et al., 2015]. Unlike most other neuromorphic systems, the SpiNNaker [Furber et al., 2014] neuromorphic supercomputer is fully programmable and combines large on-chip and external memories, distributed across the system, which enables real-time simulation of large-scale models [Rhodes et al., 2020]. This is promising for the future but, due to its prototype nature, the availability of SpiNNaker hardware is limited and even moderately-sized simulations still require a physically large system (9 boards for a model with around 100 × 10^3^ neurons and 300 × 10^6^ synapses [Rhodes et al., 2020]).

Modern GPUs have relatively little on-chip memory and, instead, dedicate the majority of their silicon area to arithmetic logic units. GPUs use dedicated hardware to rapidly switch between tasks so that the latency of accessing external memory can be ‘hidden’ behind computation, as long as there is sufficient computation to be performed. For example, the memory latency of a typical modern GPU can be completely hidden if each CUDA core performs approximately 10 arithmetic operations per byte of data accessed from memory. Unfortunately, propagating a spike in a spiking neural network simulation is likely to require accessing around 8 B of memory but perform many fewer than the required 80 instructions. This makes spike propagation highly memory bound. Nonetheless, we have shown in previous work [Knight and Nowotny, 2018] that, as GPUs have significantly higher total memory bandwidth than even the fastest CPU, moderately sized models of around 100 × 10^3^ neurons and 1 × 10^9^ synapses can be simulated on a single GPU with competitive speed and energy consumption. However, individual GPUs do not have enough memory to simulate larger brain models and, although small numbers of GPUs can be connected using the high-speed NVLink [NVIDIA Corporation, 2020a] interconnect, larger GPU clusters suffer from the same communication overheads as any other distributed HPC system.

In this work, we present a novel approach that uses the large amount of computational power available on a GPU to reduce both memory and memory bandwidth requirements and enable large-scale brain simulations on a single GPU workstation.

## 2. Results

In the following subsections, we first present two recent innovations in our GeNN simulator [Yavuz et al., 2016] which enable simulations of very large models on a GPU. We then demonstrate the power of the new features by simulating a recent model of the Macaque visual cortex [Schmidt et al., 2018b] with 4.13 × 10^6^ neurons and 24.2 × 10^9^ synapses.

### 2.1. Procedural connectivity

The first crucial innovation that enables large-scale simulations on a GPU is what we call ‘procedural connectivity’. In a brain simulation, neurons and synapses can be described by a variety of mathematical models but these are eventually all translated into time or event-driven update algorithms [Brette et al., 2007]. Our GeNN simulator [Yavuz et al., 2016] uses code generation to convert neuron and synapse update algorithms – described using ‘snippets’ of C-like code – into CUDA code for efficient GPU simulation. Before a simulation can be run, its parameters, in particular the state variables and the synaptic connectivity, need to be initialised. Traditionally, this is done by running initialisation algorithms on the main CPU prior to the simulation. The results are stored in CPU memory, uploaded to GPU memory and then used during the simulation. We have recently extended GeNN to use code generation from code snippets to also generate efficient, parallel code for model initialisation [Knight and Nowotny, 2018]. Offloading initialisation to the GPU in this way made it around 20× faster on a desktop PC [Knight and Nowotny, 2018], demonstrating that initialisation algorithms are well-suited for GPU acceleration. Here, we are going one step further. We realised that, if each synaptic connection can be re-initialised in less than the 80 operations required to hide the latency incurred when fetching its 8 B of state from memory, it could be faster and vastly more memory efficient to regenerate synaptic connections on demand rather than storing them in memory. This is the concept of procedural connectivity. It is applicable whenever synapses are static – plastic synapses which change their weights during a simulation will have to be simulated in the traditional way. Although a similar approach was used by Eugene Izhikevich for simulating an extremely large thalamo-cortical model with 1 × 10^11^ neurons and 1 × 10^15^ synapses on a modest PC cluster in 2005 [Izhikevich, 2005] – an incredible achievement – it has not been subsequently applied to modern hardware.

We implemented procedural connectivity in GeNN by repurposing our previously developed parallel initialisation methods. Instead of running them once for all synapses at the beginning of the simulation, we rerun the methods during the simulation to regenerate the outgoing synapses of each neuron that fires a spike and immediately use the identified connections and weights to run the post-synaptic code which calculates the effect of the spike onto other neurons. This is possible because the outgoing synaptic connections from each neuron are typically largely independent from those of other neurons as we shall see from typical examples below.

In the absence of knowledge of the exact microscopic connectivity in the brain, there are a number of typical connectivity schemes that are used in brain models. We will now discuss two typical examples and how they can be implemented efficiently on a GPU. One very common connectivity scheme is the ‘fixed probability connector’ for which each neuron in the presynaptic population is connected to each neuron in the postsynaptic population with fixed probability *P*_conn_. The postsynaptic targets of any presynaptic neuron can hence be sampled from a Bernoulli process with success probability *P*_conn_. One simple way of sampling from the Bernoulli process is to repeatedly draw samples from the uniform distribution Unif[0, 1] and generate a synapse if the sample is less than *P*_conn_. However, for sparse connectivity (*P*_conn_ ≪ 1), it is much more efficient to sample from the geometric distribution Geom[*P*_conn_] which governs the number of Bernoulli trials until the next success (i.e. a synapse). The geometric distribution can be sampled in constant time by inverting the cumulative density function of the equivalent continuous distribution (the exponential distribution) to obtain 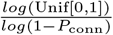 [Devroye, 2013, p499]. Note that, if we were to directly draw from the uniform distribution, the sampling for each potential synapse would be independent from any other potential synapse and all these operations could be performed in parallel. However, for the more efficient ‘geometric sampling’ employed here, the sampling for the post-synaptic targets of a presynaptic neuron must be done serially, but is still independent from the sampling for any other presynaptic neuron.

Another common scheme for defining connectivity is the ‘fixed number total connector’ in which a fixed total number *N*_syn_ of synapses is placed between randomly chosen partners from the pre- and postsynaptic populations. In order to initialise this connectivity in parallel, the number of synapses that originate from each of the *N*_pre_ presynaptic neurons must first be calculated by sampling from the multinomial distribution Mult[*N*_syn_, {*P_n_*, *P_n_*,…,*P_n_*}], where 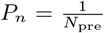, on the host CPU up front because these numbers need to add to *N*_syn_ and are hence not independent. However, once the numbers of outgoing synapses are determined, the postsynaptic targets for a presynaptic neuron can be generated very efficiently in parallel by sampling from the discrete uniform distribution Unif[0, *N*_post_] where *N*_post_ is the size of the postsynaptic population. Note, that this can only be done because the targets of each presynaptic neuron are independent from those of any other presynaptic neuron. Where synaptic weights and delays are not constant across synapses, but are described by some statistical distribution, they can also be sampled independently from each other and hence in parallel.

In order to use these parallel initialisation schemes for procedural connectivity, we require reproducible pseudorandom numbers that can be generated independently for each presynaptic neuron. In principle this could be done with ‘convential’ pseudorandom number generators (PRNGs), but each presynaptic neuron would need to maintain its own PRNG state which would lead to a significant memory overhead. Instead, we use the ‘counter-based’ Philox4×32-10 PRNG [Salmon et al., 2011]. Counter-based PRNGs are designed for parallel applications and essentially consist of a pseudo-random bijective function which takes a counter as an input (for Philox4×32-10 a 128 bit number) and outputs a random number. In constrast to convential PRNGs, this means that generating the *n*^th^ random number in a stream has exactly the same cost as generating the ‘next’ random number, allowing us to trivially divide up the random number stream between multiple parallel processes (in this case presynaptic neurons).

For an initial demonstration of the performance and scalability of procedural connectivity, we simulated a network initially designed to investigate signal propagation through cortical networks [Vogels and Abbott, 2005] but subsequently widely used as a scalable benchmark [Brette et al., 2007]. The network consists of *N* integrate-and-fire neurons, partitioned into 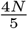 excitatory and 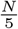 inhibitory neurons. The two populations of neurons are connected to each other and with themselves with fixed probability *P*_conn_ = 10 %.

We ran simulations of this network at scales ranging from 1 × 10^3^ to 1 × 10^6^ neurons (100 × 10^3^ to 100 × 10^9^ synapses respectively) on a representative selection of NVIDIA GPU hardware: Jetson TX2, a low-power embedded system with 8 GB (shared memory); Geforce MX130, a laptop GPU with 2 GB; Geforce GTX 1650, a low-end desktop GPU with 4 GB; and Titan RTX, a high-end workstation GPU with 24 GB. Fig. 1 shows the duration of these simulations using our new procedural approach or using the standard approach of storing synaptic connections in memory employing two different data structures. Both data structures are described in more detail in our previous work [Knight and Nowotny, 2018] but briefly, in the ‘sparse’ data structure, a presynaptic neuron’s postsynaptic targets are represented as an array of indices whereas, in the ‘bitfield’ data structure, they are represented as a *N*_post_ array of bits where a ‘1’ at position *i* indicates that there is a connection to postsynaptic neuron *i* and a ‘0’ that there is not. None of our devices have enough memory to store the 100 × 10^9^ synapses required for the largest scale using either data structure but, at the 100 × 10^3^ neuron scale, the bitfield data structure allows the model to fit into the memory of several devices it otherwise would not. However, not only is the new procedural approach the *only* way of simulating models at the largest scales but, as Fig. 1 illustrates, even at smaller scales the performance of the precedural approach is competitive with and sometimes better than the standard approach. All of the synapses in this model have the same synaptic weight meaning that they can be hard-coded into the procedural connectivity kernels. However, if weights vary across synapses, the ‘bitfield’ cannot be used and the memory constraints for the ‘sparse’ representation become even more severe.

**Figure 1.**
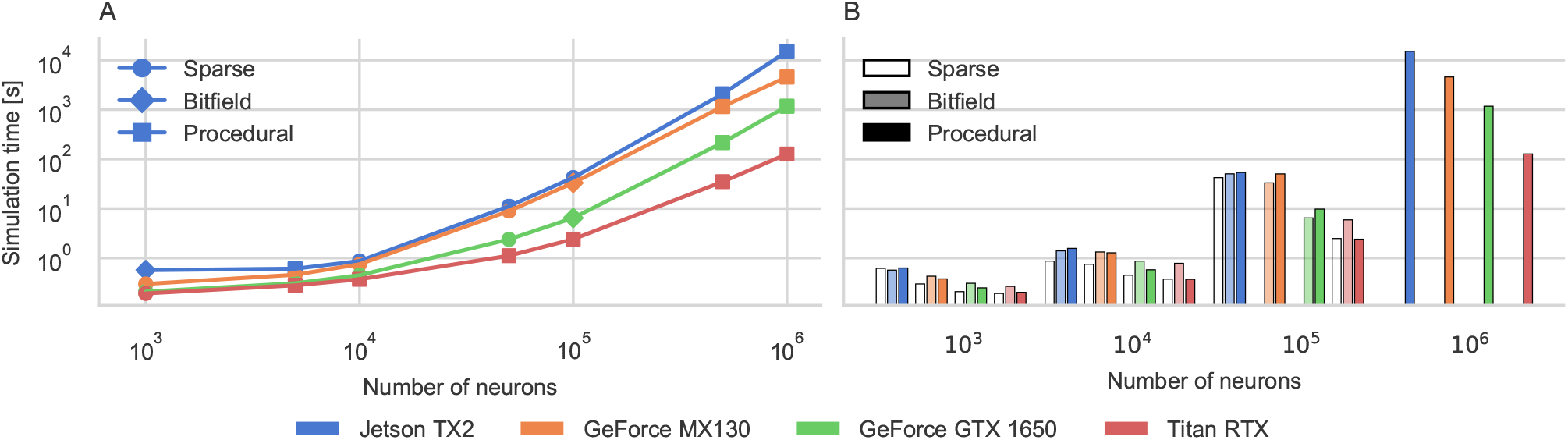
Simulation time performance scaling on a range of modern GPUs (colors). **A** The best performing approach at each scale on each GPU (indicated by the symbols). For the largest models, the procedural method is always best. **B** Raw performance of each approach on each GPU. Missing bars indicate insufficient memory to simulate.

### 2.2. Kernel merging

NVIDIA GPUs are typically programmed in CUDA using a Single Instruction Multiple Thread (SIMT) paradigm where programmers write ‘kernel’ functions containing serial C-like code which is then executed in parallel across many virtual threads. We call our second innovation ‘kernel merging’ and it relates to the way these kernels are implemented. While the procedural connectivity presented in the previous section allows simulating models which would otherwise not fit into the memory of a GPU, there are additional problems when using code generation for models with a large number of neuron and synapse populations. GeNN and other SNN simulators which use code generation to generate all of their simulation code [Blundell et al., 2018] (as opposed to, for example NESTML [Plotnikov et al., 2016], which uses code generation only to generate neuron simulation code) generate seperate pieces of code for each population of neurons and synapses. This allows optimizations such as hard-coding constant parameters and, although generating code for models with many populations will result in large code size, C++ CPU code can easily be divided between multiple modules and compiled in parallel, minimizing the effects on build time. However, GPUs can only run a small number of kernels – which are equivalent to modules in this context – simultaneously (128 on the latest NVIDIA GPUs [NVIDIA Corporation, 2019, p278]). Therefore, in GeNN, multiple neuron populations are simulated within each kernel, resulting in code of the form shown in the following pseudocode for simulating 3 populations of 100 neurons each in a single kernel:

~~~
**void** updateNeurons () {
**if**(thread < 100) {
*// Update neuron population A*
} **else if**(thread >= 100 && thread < 200) {
*// Update neuron population B*
} **else if**(thread >= 200 && thread < 300) {
*// Update neuron population C*
}
}
~~~

This works well for a small number of populations but, as Fig. 2A illustrates, when we partition a model consisting of 1 000 000 LIF neurons into an increasingly large number of (smaller and smaller) populations, compilation time increases super-linearly and quickly becomes impractical. Furthermore, the simulation also runs more slower with a large number of populations (Fig. 2B). Normally, we would expect this model to be memory bound as each thread in the model reads 32 B of data and, as discussed above, hiding the latency of these memory accesses would require approximately 320 arithmetic operations – many more than required to sample an input current from the normal distribution and update a LIF neuron. Fig. 2C – obtained using data from the NVIDIA Nsight compute profiler [NVIDIA Corporation, 2020b] – shows that this is true for small numbers of populations. In this case, the memory system is around 90 % utilised. However, when the model is partitioned into larger numbers of smaller populations, the memory is used less efficiently and the kernel becomes latency bound, i.e. neither memory *nor* compute are used efficiently. Investigating further, we found that this drop in performance was accompanied by an increasing number of “No instruction” stalls (Fig. 2D) which are events that prevent the GPU from doing any work during a clock cycle. These particular events are likely to be caused by “Excessively jumping across large blocks of assembly code” [NVIDIA Corporation, 2020b, p47], which makes sense as we are generating kernels with hundreds of thousands of lines of code. Several neural modelling tools including Brian2 [Stimberg et al., 2014] provide modellers with tools to work with ‘slices’ of neuron populations, allowing models to be defined with fewer populations. However, if a model is defined by connecting these slices together, the resulting connectivity is the result of *multiple* simple connection rules of the type discussed in the previous section, making it much more difficult to apply our procedural connectivity approach. Furthermore, such an approach places the responsibility for structuring a model in such a way that it can be simulated efficiently onto the modellers, who often prefer to concentrate on the science and organise populations according to anatomy or physiology.

**Figure 2.**
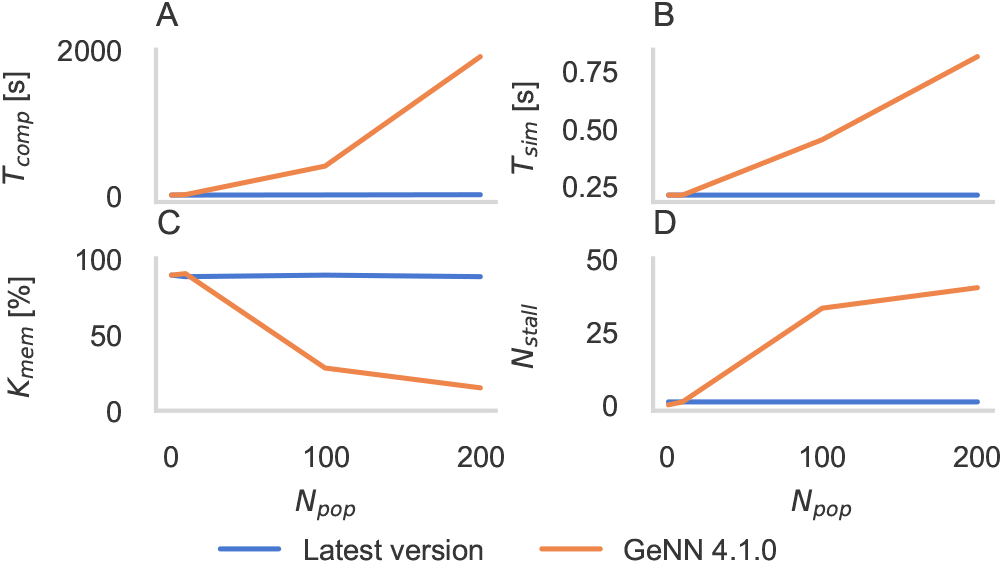
Performance of a simulation of 1 000 000 LIF neurons driven by a gaussian input current, partitioned into varying numbers (*N_pop_*) of populations and running on a workstation equipped with a Titan RTX GPU. **A** Compilation time (*T_comp_*) using GCC 7.5.0. **B** Simulation time (*T_sim_*) for an 1 s simulation. **C** Memory throughput (*K_mem_*) reported by NVIDIA Nsight compute profiler “Speed of light” metric. **D** Number of “No instruction” stalls reported by NVIDIA Nsight compute profiler (*N_stall_*).

To address the issue of too many populations, we developed a new code generator for GeNN which first ‘merges’ the model description, grouping together populations which can be simulated using the same generated code. From this merged description, structures are generated to store the pointers to state variables and parameter values which are still allowed to differ between merged populations:

~~~
**struct** NeuronUpdateGroup {
**unsigned int** numNeurons;
**float**∗ V;
};
~~~

An array of these structures is then declared for each merged population and each element is initialised with pointers to state variables and parameter values:

~~~
NeuronUpdateGroup neuronUpdateGroup [3];
neuronUpdateGroup [0] = {100, VA};
neuronUpdateGroup [1] = {100, VB};
neuronUpdateGroup [2] = {100, VC};
~~~

where VA is a pointer to the array containing the state variable ‘V’ of populations ‘A’ and so on. In order for a thread to determine which neuron in which population it should simulate, we generate an additional data structure – an array containing a cumulative sum of threads used for each population:

~~~
**unsigned int** start Thread [3] = {0, 100, 200};
~~~

Each thread performs a simple binary search within this array to find the index of the neuron and population it should simulate. As Fig. 2 shows, this approach solves the observed issues with compilation time and simulation performance.

### 2.3. The multi-area model

Due to lack of computing power and sufficiently detailed connectivity data, previous models of the cortex have either focussed on modelling individual local microcircuits at the level of individual cells [Izhikevich and Edelman, 2008, Potjans and Diesmann, 2014] or modelling multiple connected areas at a higher level of abstraction [Cabral et al., 2014]. However, recent data [Belitski et al., 2008] has shown that cortical activity has distinct features at both the global and local levels which can only be captured by modelling interconnected microcircuits at the level of individual cells. The recent multi-area model [Schmidt et al., 2018a,b] is an example of such multi-scale modeling. It uses scaled versions of a previous, 4 layer microcircuit model [Potjans and Diesmann, 2014] to implement 1 mm^2^ ‘patches’ for 32 areas of the macaque visual cortex. The patches are connected together according to inter-area axon tracing data from the CoCoMac [Bakker et al., 2012] database, further refined using additional anatomical data [Markov et al., 2014a] and heuristics [Ercsey-Ravasz et al., 2013] to obtain estimates for the number of synapses between areas. The synapses are distributed between populations in the source and target area using layer-specific tracing data [Markov et al., 2014b] and cell-type-specific dendritic densities [Binzegger et al., 2004]. Individual populations are connected by the fixed number connectors described above. For a full description of the multi-area model please see Schmidt et al. [2018a,b]. In 2018, this model was simulated using NEST [Gewaltig and Diesmann, 2007] on one rack of an IBM Blue Gene/Q supercomputer (a 2 m high enclosure containing 1024 compute nodes, weighing over 2 t and requiring around 80 kW of power). On this system, initialization of the model took around 5 min and simulating 1 s of biological time took approximately 12 min [Schmidt et al., 2018b].

The multi-area model consists of 4.13 × 10^6^ neurons in 254 populations and 24.2 × 10^9^ synapses in 64 516 populations. Without kernel merging, it would therefore be unlikely that the model would compile or simulate at a workable speed using GeNN. Additionally, unlike the model we benchmarked previously, each synapse in this model has an independant weight and synaptic delay sampled from a normal distribution so the bitfield data structure cannot be used. Even if we assume that 16 bit floating-point would provide sufficient weight precision, that delays could be expressed as 8 bit integers and that neuron populations are all small enough to be indexed using 16 bit indices, our sparse data structure would still require 5 B per synapse, such that the complete synaptic data would need over 100 GB of GPU memory. While a cluster of GPUs connected using NVLink could be built with this much memory, it is more than any single GPU has available. However, using procedural connectivity, we are able to simulate this model on a single workstation with a Titan RTX GPU.

In order to validate our GeNN simulations, we ran a 10.5 s simulation of the multi-area model in a ‘ground state’ where inter-area connections have the same strength as intra-area connections and a 100.5 s simulation in a ‘resting state’ where inter-area connections are 1.9× stronger. Initialization of our model took 6 min (3 min of which was spent generating and compiling code) and simulation of each biological second took 7.7 min in the ground state and 8.4 min in the resting state– 35 % and 30 % less than the supercomputer simulation respectively. Fig. 3A-C shows some example spike rasters from three of the modelled areas, illustrating the asynchronous irregular nature of the model’s ground state whereas Fig. 3D-F illustrate the characteristic irregular activity and population bursts of the same areas in the resting state. Next, we calculated the per-layer distributions of rates, spike-train irregularity and cross-correlation coefficients across all areas (disregarding the first 500 ms of simulation) and compared them to the same measures obtained from spike trains generated by the supercomputer simulations. We calculated irregularity using the revised local variation LvR [Shinomoto et al., 2009], averaged over a subsample of 2000 neurons and cross-correlation from spike histograms with 1 ms bins, calculated from a subset of 2000 non-silent neurons. The violin plots in Fig. 3G-L show the comparison of the distributions of values obtained from the NEST and GeNN simulations in both states – which are essentially identical.

**Figure 3.**
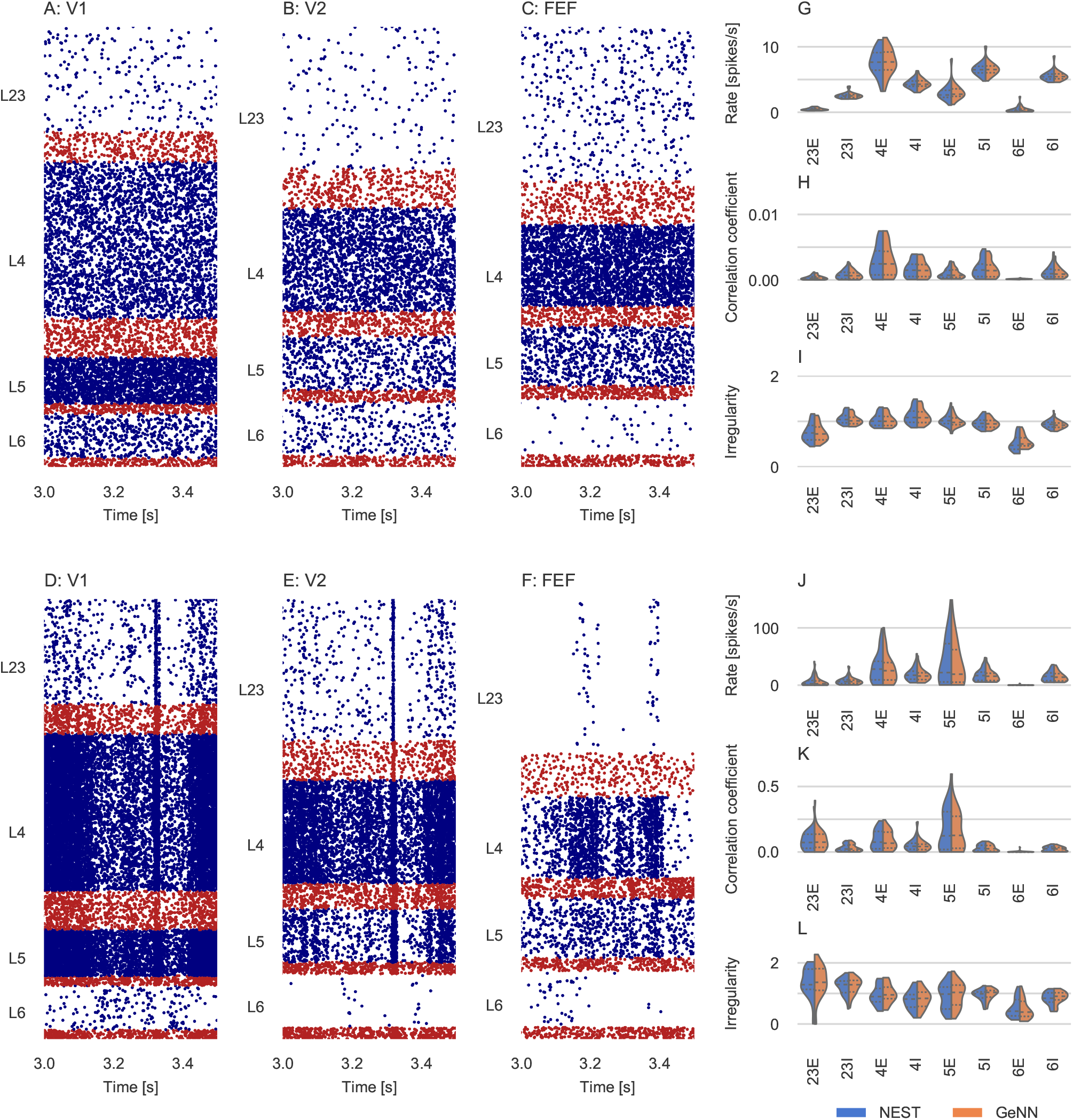
Results of full-scale multi-area model simulation in ground and resting states. **A-F** Raster plots of spiking activity of 3 % of the neurons in area V1 (A,D), V2 (B,E), and FEF (C,F). Blue: excitatory neurons, red: inhibitory neurons. **G-L** Spiking statistics for each population across all 32 areas simulated using GeNN and NEST shown as split violin plots. Solid lines: medians, Dashed lines: Interquartile range. **G,J** Population-averaged firing rates. **H,K** Average pairwise correlation coefficients of spiking activity. **I,L** Irregularity measured by revised local variation LvR [Shinomoto et al., 2009] averaged across neurons.

## 3. Discussion

In this work we have presented a novel approach for large-scale brain simulation on GPU devices which entirely removes the need to store connectivity data in memory. We have shown that this approach allows us to simulate a cortical model with 4.13 × 10^6^ neurons and 24.2 × 10 synapses [Schmidt et al., 2018a,b] on a single modern GPU. While this represents a significant step forward in terms of making truly large-scale brain modelling tools accesible to a large community of brain researchers, this model still has around 20× fewer neurons and 40× fewer synapses than the brain of even a small mammal such as a mouse [Herculano-Houzel et al., 2006]. Our implementation of the multi-area model requires a little over 12 GB of GPU memory, with the majority (8.5 GB) being used for the circular dendritic delay buffers (see Knight and Nowotny [2018]). These are a per-neuron (rather than per-synapse) data structure but, because the inter-area connections in the model have delays of up to 500 simulation timesteps (0.1 ms), the delay buffers become quite large.

One important aspect of large-scale brain simulations not addressed in this work is synaptic plasticity and its role in learning. As discussed by Knight and Nowotny [2018], GeNN supports a wide variety of synaptic plasticity rules. In order to modify synaptic weights, they need to be stored in memory rather than generated procedurally. However, connectivity could still be generated procedurally, potentially halving the memory requirements of models with synaptic plasticity. This would be sufficient for synaptic plasticity rules that only require access to presynaptic spikes and postsynaptic neuron states Brader et al. [2007], Clopath et al. [2010] but, for many Spike-Timing-Dependent Plasticity (STDP) rules, access to *postsynaptic* spikes is also required. GeNN supports such rules by automatically generating a lookup table structure (see Knight and Nowotny [2018]). While this process could be adapted to generate a lookup table from procedural connectivity, this would further erode memory savings. However, typically not all synapses in a simulation are plastic and those that are not could be simulated fully procedurally.

In this work, we have discussed the idea of procedural connectivity in the context of GPU hardware but, we believe that there is also potential for developing new types of neuromorphic hardware built from the ground up for procedural connectivity. Key components such as the random number generator could be implemented directly in hardware leading to truly game-changing compute time improvements.

## 4. Methods

In all experiments presented in this work, neurons are modelled as leaky integrate-and-fire (LIF) units with the parameters listed in Table 1. The membrane voltage *V_i_* of neuron *i* is modelled as

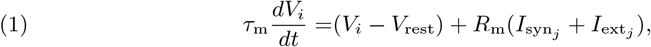

where *τ*_m_ and *R*_m_ represent the time constant and resistance of the neuron’s cell membrane, *V*_rest_ defines the resting potential, 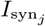 represents the synaptic input current and 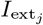 represents an external input current. When the membrane voltage crosses a threshold *V*_th_ a spike is emitted, the membrane voltage is reset to *V*_rest_ and updating of *V* is suspended for a refractory period *τ*_ref_. In the models where there are synaptic connections, pre-synaptic spikes lead to exponentially-decaying input currents 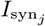

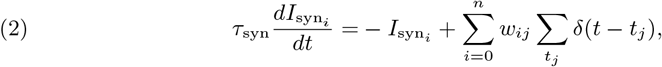

where *τ*_syn_ represents the decay time constant and *t_j_* are the arrival times of incoming spikes from *n* presynaptic neurons. The continuous terms of the Eq. 1 and 2 are seperately solved algebraically so that the synaptic input current 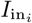 is treated as a constant throughout each simulation timestep.

**Table 1.** Model parameters.

| Parameter | Procedural connectivity<br>benchmark | Merging<br>benchmark | Multi-area<br>model |
| --- | --- | --- | --- |
| $\tau_m$ [ms] | 20 | 20 | 2 |
| $V_{\text{rest}}$ [mV] | -60.0 | -70.0 | -65 |
| $V_{\text{th}}$ [mV] | -50.0 | -51.0 | -50 |
| $R_m$ [MΩ] | 20 | 20 | 40 |
| $\tau_{\text{syn}}$ [ms] | 5/10<sup>1</sup> | — | 0.5 |
| $\tau_{\text{ref}}$ [ms] | 5 | 2 | 2 |
| $I_{\text{ext}_j}$ [nA] | 0.55 | 1.00 ± 0.25 | Poisson<sup>2</sup> |
| $w_{ij}$ [nA] | $\frac{3.2}{N} / \frac{40.8}{N}$<sup>1</sup> | — | Various<sup>2</sup> |

<sup>1</sup>Excitatory/Inhibitory.<sup>2</sup>Please refer to Schmidt et al. [2018b, Table 1,2]

## 5. Acknowledgements

We would like to thank Jari Pronold, Sacha van Albada, Agnes Korcsak-Gorzo and Maximilian Schmidt for their help with the multi-area model data; and Dan Goodman and Mantas Mikaitis for their feedback on the manuscript. This work was funded by the EPSRC (Brains on Board project, grant number EP/P006094/1).

## 6. Author contributions

J.K. and T.N. wrote the paper. T.N. is the original developer of GeNN. J.K. is currently the primary GeNN developer and was responsible for extending the code generation approach to the procedural simulation of synaptic connectivity. J.K. performed the experiments and the analysis of the results that are presented in this work.

